# Cost of preterm birth during initial hospitalization: A care provider’s perspective

**DOI:** 10.1101/532713

**Authors:** Hadzri Zainal, Maznah Dahlui, Shahrul Aiman Soelar, Tin Tin Su

## Abstract

Preterm birth incidence has risen globally and remains a major cause of neonatal mortality despite improved survival. The demand and cost of initial hospitalization has also increased. This study assessed care provider cost in neonatal intensive care units of two hospitals in the state of Kedah, Malaysia. It utilized universal sampling and prospectively followed up preterm infants till discharge. Care provider cost was assessed using mixed method of top down approach and activity based costing. A total of 112 preterm infants were recruited from intensive care (93 infants) and minimal care (19 infants). Majority were from the moderate (23%) and late (36%) preterm groups followed by very preterm (32%) and extreme preterm (9%). Mean total cost per infant increased with level of care and degree of prematurity from MYR 2,751 (MYR 374 - MYR 10,103) for preterm minimal care, MYR 8,478 (MYR 817 - MYR 47,354) for late preterm intensive care to MYR 41,598 (MYR 25,351- MYR 58,828) for extreme preterm intensive care. Mean cost per infant per day increased from MYR 401 (MYR 363- MYR 534), MYR 444 (MYR 354 – MYR 916) to MYR 532 (MYR 443-MYR 939) respectively. Cost was dominated by overhead (fixed) costs for general (hospital), intermediate (clinical support services) and final (NICU) cost centers where it constituted at least three quarters of mean admission cost per infant while the remainder was consumables (variable) costs. Breakdown of overhead cost showed NICU specific overhead contributing at least two thirds of mean admission cost per infant. Personnel salary made up three quarters of NICU specific overhead. Laboratory investigation was the cost driver for consumables ranging from 29% (intensive care) to 84% (minimal care) of mean total consumables cost per infant. Gender, birth weight and length of stay were significant factors and cost prediction was developed with these variables.

## INTRODUCTION

Preterm birth is defined as delivery before 37 completed weeks of gestation. It can be categorized into late preterm (34 weeks to less than 37 weeks gestation), moderate preterm (32 weeks to less than 34 weeks gestation), very preterm (28 weeks to less than 32 weeks gestation) and extremely preterm (less than 28 weeks gestation) (1). However most morbidity and mortality affect very preterm and extremely preterm infants (2). Preterm birth is increasingly common with substantial medical, economic and social impact as it is invariably associated with acute and chronic complications (3, 4). Due to advancements in care over the last few decades, outcome and survival of preterm infants have improved, however, the economic impact of preterm care has gained much attention. Economic evaluation on the cost of managing preterm infants can generally be divided into intensive care costs during initial hospitalization and long term costs such as health and educational needs during the early years. Most studies have been devoted to costs of intensive care as initial hospitalization accounts for the bulk of health care cost during the first 2 years of life of a preterm infant (5). More specifically, costs during initial hospitalization dominate 92.0% of the incremental costs per preterm survivor (6). Degree of prematurity also affects the cost of managing preterm infants. Although moderate preterm infants have much less complications and better survival rate, substantial resources are still needed to manage them as they comprise the bulk of preterm admissions. On the other hand, very preterm and extremely preterm infants may be less in number but they require intense care and longer hospital stay (7–9).

Since its inception in 2009, Malaysia’s preterm birth registry showed an increasing rate from 8.1% to 11.3% between 2010 and 2012 (10). Its neonatal intensive care service is largely public funded and run in more than 38 government hospitals (11, 12). In terms of workload preterm infants made up more than 60% of all neonatal intensive care unit neonatal intensive care unit (NICU) admissions and more than 60% of babies below 1500g were ventilated in NICUs of government hospitals throughout Malaysia with mean ventilation days of 6.6 days. Malaysia currently relies on studies from abroad for economic burden of preterm birth. There has been only one such study locally which found NICU services for infants between 1000 and 1500 g birth weight to be cost effective (11). As money is a limited resource for the care provider an economic assessment is vital for greater efficiency of care. Findings from this study may aid neonatal care policy planning and services for optimal management and improved outcome of preterm infants.

## METHODOLOGY

### Study design and participants

This cost of illness study utilized universal sampling and prospectively followed up preterm infants from admission till discharge during initial hospitalization. Study was conducted at the NICUs of Hospital Sultanah Bahiyah (Center 1) and Hospital Sultan Abdul Halim (Center 2) the largest hospitals and referral centres for the state of Kedah which has one of the highest mean number of neonatal admissions per hospital in Malaysia (12). Center 1 offered tertiary level care and had an average of 800 preterm admissions annually while Center 2 had secondary level care with an average of 500 preterm admissions annually. Inborn and out born preterm infants delivered via normal delivery or Caesarean section and admitted to NICU of both centers during data collection period were included in this study. Excluded were preterm infants admitted for less than 24 hours (for observation) and preterm infants with severe congenital anomalies as these infants would only receive supportive care due to their often short duration of life. Total of 101 preterm infants consecutively admitted to intensive and intermediate care and 20 preterm infants admitted to minimal care were recruited. However 6 preterm infants were excluded with reasons leaving 112 preterm infants in the final group for analysis. Data was collected over a period of six months from January 1st to June 30th 2015.

### Cost data collection

Care provider costing was conducted from the perspectives of the hospital, NICU and supporting clinical services using mixed method of top down approach (fixed overhead cost) and activity based costing (variable consumables cost) to obtain total admission cost per preterm infant by degree of prematurity (Table 1). Care provider cost was assessed based on the NICU preterm pathway of care which was generally similar at the study centers. For cost prediction, independent variables included were hospital (NICU) of admission, length of stay, ventilation duration, gender, gestation and birth weight while outcome measured was total admission cost per preterm infant.

**Table 1:**
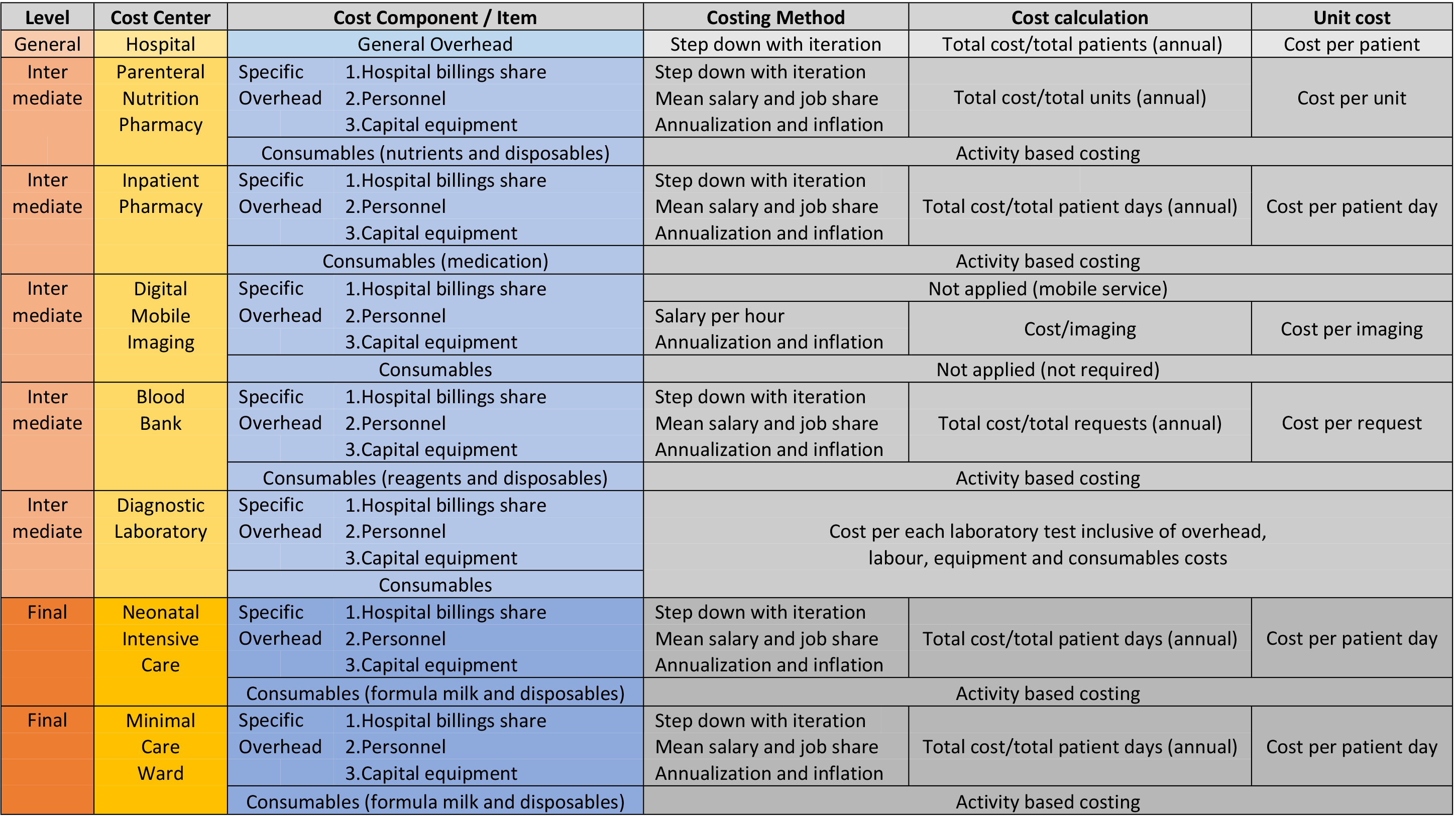
Care provider cost analysis framework

Overhead cost was estimated using top-down approach. It included general hospital overhead and specific overhead for each intermediate (clinical support services) and final (NICU) cost centers. Administration and non-medical staff salaries, utilities (electricity, water and telecommunication services), non-clinical hospital support service (i.e. cleaning, laundry, pest control, maintenance and repairs), hospital information system and security services constituted general hospital overhead. These were expenses associated with running of the hospital and shared among patients who utilized hospital in-patient and out-patient services. Share of hospital billings for utilities and non-clinical support services (based on floor area ratio), personnel salary and annual equipment costs made up specific overhead for each intermediate and final cost centers. Data for calculation of general and specific cost center overheads (step down with iteration) such as personnel, equipment, utilities and billings costs were obtained retrospectively from 2015 hospital records. Preterm infants enrolled were prospectively followed up from admission till discharge and consumables cost for each preterm infant was calculated using activity based costing through observation of resource consumption. All expenditure pertaining to this was determined, valued and subsequently unit cost for each level of activity was calculated for each patient. Consumable items included disposables, medication, enteral and parenteral nutrition, infusion, transfusion, blood investigation and imaging. Consumables calculated include those used by intermediate cost centers for management of preterm infants enrolled in this study. Mean total admission cost per preterm infant was determined by summing up total overhead cost per infant for general, intermediate and final cost centers with total consumables cost per infant. Data recording tool also documented information such as patient identification, gestation, birth weight, dates of admission and discharge and number of days ventilated (invasive and non-invasive).

### Statistical analysis

All cost data were presented in Malaysian Ringgit (MYR) where $1 = MYR 3.75 (prevailing conversation rate in 2015). Cost data were tabulated with Microsoft Excel (2010) and subsequently analyzed with IBM SPSS Statistics version 22. Descriptive analysis such as means with range for continuous variables and frequencies and percentages for the categorical variables were used. Provider cost was calculated by degree of prematurity for mean total cost per infant, mean cost per patient day, mean cost per infant by cost components and mean cost per infant by items for consumables. Multifactorial ANOVA was used to identify independent variables with statistically significant adjusted means. These variables were then included for cost prediction using general linear model. Regression coefficients and power of regression were obtained. Significance level was set at p <0.05.

## RESULTS

### Infant characteristics

101 preterm infants from intensive care and 20 preterm infants from minimal care (moderate and late preterm) were recruited from both study centres. 6 mortalities from the intensive care group and 3 preterm infant with incomplete data from the minimal care group were excluded. This left a total of 112 preterm infants for analysis from both intensive care (93 infants) and minimal care (19 infants) groups (Table 2). Majority of preterm infants included were from the moderate (23%) and late preterm groups (36%) followed by very preterm (32%) and extreme preterm (9%). In intensive care extreme preterm group had the lowest mean birth weight of 0.92kg (0.61kg-1.26kg) but longest mean duration of admission at 84 days (27 days – 132 days) and mean ventilation days that extended up to 53 days (22 days-104 days). In contrast late preterm had the highest mean birth weight of 2.08kg (0.81kg-2.70kg) but shortest mean hospital stay at 19 days (2 days – 104 days) and lowest mean ventilation duration of 4 days (0 days-20 days). Preterm infants in minimal care had the shortest mean hospital stay of 7 days (1 days – 27 days) while the single longest hospital stay was 167 days for an infant in the moderate preterm group.

**Table 2:** Preterm characteristics

| Characteristics | Center 1 and Center 2 (n=112) |  |  |  |  |
| --- | --- | --- | --- | --- | --- |
|  | Mean (range) |  |  |  |  |
|  | Minimal<br>Care<br>n=19 | Intensive Care |  |  |  |
|  |  | Late preterm<br>n=23 | Moderate preterm<br>n=24 | Very preterm<br>n=36 | Extreme preterm<br>n=10 |
| Birth weight<br>(kg) | 2.06<br>(1.44-2.80) | 2.08<br>(0.81-2.70) | 1.69<br>(0.99-2.73) | 1.26<br>(0.88-2.25) | 0.92<br>(0.61-1.26) |
| Length of stay<br>(days) | 7<br>(1-27) | 19<br>(2-104) | 33<br>(7-167) | 51<br>(9-94) | 84<br>(27-132) |
| Ventilation<br>(days) | -<br>- | 4<br>(0-20) | 6<br>(1-43) | 16<br>(1-68) | 53<br>(22-104) |

### Cost analysis

Mean total cost per infant increased with level of care and prematurity from MYR 2,751 (MYR 374 - MYR 10,103) for preterm minimal care, MYR 8,478 (MYR 817 - MYR 47,354) for late preterm intensive care, MYR 16,557 (MYR 2,643 – MYR 98,249) for moderate preterm intensive care, MYR 24,555 (MYR 3,648 – MYR 48,943) for very preterm intensive care and MYR 41,598 (MYR 25,351-MYR 58,828) for extreme preterm intensive care (Table 3). Mean cost per infant per day similarly increased from MYR 401 (MYR 363-MYR 534), MYR 444 (MYR 354 – MYR 916), MYR 463(MYR 351 – MYR 619), MYR486 (MYR 376 - MYR 782) and MYR 532 (MYR 443-MYR 939) respectively. Overhead cost included general hospital overhead and specific overhead for each intermediate (clinical support services) and final (NICU) cost centres. Mean total overhead cost per infant ranged from MYR 2,518 (MYR 362-MYR 9,406) in minimal care to MYR 31,527 (MYR 12,710-MYR 47,268) in extreme preterm intensive care. Breakdown of overhead cost revealed NICU specific overhead as the overwhelming contributor in all categories (69% - 89%). Mean total consumables cost per infant ranged from MYR 232 (MYR 12- MYR 697) in minimal care to MYR 10,071 (MYR 7,428- MYR 13,441) in extreme preterm intensive care.

**Table 3:** Cost per preterm admission

| Cost component | Mean (range) MYR |  |  |  |  |
| --- | --- | --- | --- | --- | --- |
|  | Minimal care<br>*n*=19 | Intensive care |  |  |  |
|  |  | Late preterm<br>*n*=23 | Moderate preterm<br>*n*=24 | Very preterm<br>*n*=36 | Extreme preterm<br>*n*=10 |
| General | 16 | 16 | 16 | 16 | 16 |
| overhead* | - | - | - | - | - |
| Parenteral nutrition | 0 | 190 (2%) | 975 (6%) | 1296 (5%) | 2028 (5%) |
| pharmacy overhead* | 0 | (0-2730) | (0-8190) | (0-4368) | (624-3120) |
| Inpatient pharmacy | 43 | 113 | 196 | 308 | 505 |
| overhead* | (6-162) | (12-624) | (42-1002) | (54-564) | (162-792) |
| Mobile imaging | 8 | 83 | 109 | 133 | 269 |
| overhead* | (0-48) | (24-384) | (24-552) | (0-384) | (120-432) |
| Blood bank | 0 | 2 | 14 | 21 | 81 |
| overhead* | 0 | (0-28) | (0-196) | (0-112) | (28-140) |
| NICU | 2452 (89%) | 6401 (75%) | 11107 (67%) | 17444 (71%) | 28628 (69%) |
| overhead* | (340-9180) | (680-35360) | (2380-56780) | (3060-31960) | (9180-44880) |
| Total | 2518 | 6806 | 12417 | 19217 | 31527 |
| overhead | (362-9406) | (732-39046) | (2462-66688) | (3154-36600) | (12710-47268) |
| Consumables | 232 (8%) | 1673 (20%) | 4159 (25%) | 5338 (22%) | 10071 (24%) |
| cost | (12-697) | (85-9243) | (181-31561) | (494-13393) | (7428-13441) |
| Total admission | 2751 (100%) | 8478 (100%) | 16557 (100%) | 24555 (100%) | 41598 (100%) |
| cost | (374-10103) | (817-47354) | (2643-98249) | (3648-48943) | (25351-58828) |

\* Overhead cost items \$1 = MYR 3.67 Percentages shown for main cost contributors

Further scrutiny showed personnel cost as the cost driver (76%) of NICU overhead. This was followed by annual equipment cost (13%) while utilities and auxiliary services collectively made up just a little more than 10% of total NICU specific overhead. Mean total consumables cost per infant increased with level of care and prematurity and this pattern replicated throughout mean costs for individual consumable items. For all levels of care and preterm categories laboratory investigation was the cost driver for consumables ranging from 29% (intensive care) to 84% (minimal care) of mean total consumables cost. Other major contributors were medication, parenteral nutrition and disposables. Parenteral nutrition, transfusion and disposables costs were especially higher in the extreme preterm group making up 18%, 17% and 21% respectively of total consumables cost

### Cost prediction

To obtain the cost predictors multifactorial ANOVA was performed using variables of hospital (NICU) of admission, length of stay, ventilation duration, gender, gestation and birth weight. From the adjusted means, gender, length of stay and birth weight were the significant factors. Total admission cost per infant was higher with increased length of stay, reduced birth weight and the male gender. Subsequently non-significant variables were removed and multifactorial ANOVA was repeated with length of stay, gender and birth weight to improve the cost prediction and observed power (Table 4). This yielded a general linear regression equation of:

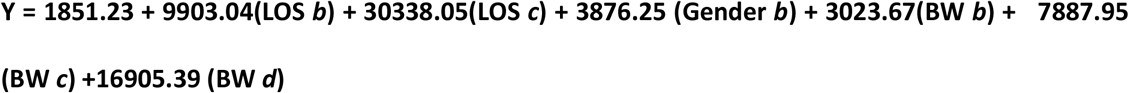

Whereby:

Y = total admission cost per infant;
LOS *i* = length of stay; LOS ref = less than 1 month; LOS *b* = 1 to 2 month; LOS *c* = more than 2 months Gender ref = female; Gender *b* = male
BW *i* = birth weight; BW ref = 2kg and more; BW *b* = 1.5 to 1.99 kg; BW *c* = 1.0 to 1.49 kg; BW *d* = less than 1 kg

**Table 4:** Final variables for cost prediction (multifactorial ANOVA)

| Variables | Coefficient<br>B | SE | Total admission cost |  | Multiple<br>Comparisons | p-value | Observed<br>Power |
| --- | --- | --- | --- | --- | --- | --- | --- |
|  |  |  | Mean<br>(MYR) | SE<br>(MYR) |  |  |  |
| Intercept | 1851.23 | 1385.86 |  |  |  |  |  |
| **Length of stay** |  |  |  |  |  | **<0.05** | 1.000 |
| A. < 1month | Ref | - | 10743.61 | 1315.90 | A vs B | <0.05 |  |
| B. 1-2 month | 9903.04 | 2107.53 | 20646.65 | 1460.81 | A vs C | <0.05 |  |
| C. >2 months | 30338.05 | 2774.55 | 41081.66 | 1969.68 | B vs C | <0.05 |  |
| **Gender** |  |  |  |  |  | **<0.05** | 0.803 |
| A. Male | 3876.25 | 1364.86 | 26095.43 | 1129.48 | - | - |  |
| B. Female | Ref |  | 22219.18 | 989.18 | - | - |  |
| **Birth weight** |  |  |  |  |  | **<0.05** | 0.997 |
| A. ≥2kg | Ref | - | 17203.05 | 1823.71 | A vs B | 1 |  |
| B. 1.5-1.99kg | 3023.67 | 1898.54 | 20226.72 | 1829.08 | A vs C | 0.035 |  |
| C. 1.0-1.49kg | 7887.95 | 2341.37 | 25091.00 | 1222.21 | A vs D | <0.05 |  |
| D. <1kg | 16905.39 | 3149.06 | 34108.44 | 2112.67 | B vs C | 0.149 |  |
|  |  |  |  |  | B vs D | <0.05 |  |
|  |  |  |  |  | C vs D | <0.05 |  |

## DISCUSSION

Apnoea of prematurity and respiratory distress syndrome are among complications that can occur with increasing prematurity. These conditions often require intensive care for ventilation, cardiovascular and nutritional support from day one of life. Other complications may ensue such as intra-ventricular haemorrhage, nosocomial pneumonia and sepsis (13). These conditions may lead to prolonged NICU stay in order to stabilize, establish feeding and gain optimal weight. Thus increasing prematurity and reduced birth weight entail intense resource utilization (9). In this study the inverse relation was demonstrated and most pronounced in the extreme preterm group which had the lowest mean birth weight but highest mean length of stay and ventilation days. The consequence is that intensity of care and duration are two parameters that influence cost of neonatal intensive care(14). Intensity is factored by quantity and cost of resources (such as ventilation) utilized per admission day while duration is represented by length of stay. This study also demonstrated the increase in mean cost per admission with level of care and degree of prematurity. This inverse relationship is compatible with literature where studies have shown that increasing prematurity equates to intense resource utilization that translates into higher cost (3–5, 14, 15). Although mean cost per day for a preterm infant increased with the level of care and prematurity in this study, increments in cost per day between levels of care and preterm categories were not pronounced (5% - 10%). This can be attributed to the fixed overhead cost that contributed to the bulk of total admission cost per infant at every level of care and preterm category (75% - 92%). This was inclusive of general hospital overhead and specific overhead for intermediate (clinical support services) and final (NICU) cost centres. Consequently, it left a reduced percentage for the variable component of consumables cost that differed with patient workload. This was consistent with a previous study which revealed that close to 80% of intensive care unit cost is actually fixed such as for personnel and equipment (16). Further analysis revealed NICU specific overhead accounting for at least two thirds of mean admission cost per infant. Personnel salary made up three quarters of NICU specific overhead. Other studies similarly found personnel cost as the cost driver for NICU preterm care (11, 15, 17).

In this study mean cost per preterm admission ranged between MYR 8,478 ($2,310) for late preterm to MYR 41,598 ($11,335) for extreme preterm. Meanwhile mean cost per day ranged from MYR 444 ($121) for late preterm to MYR 532 ($145) for extreme preterm. Comert et al found mean total cost of $4187 and mean cost per day of $303 (18). Akman et al and Geitona et al recorded mean cost per preterm of $4345 and $6,801 respectively (15, 19). Narang et al calculated mean cost per day of $125 (17). Kırkby et al reported a mean intensive care cost of $31,000 for preterm infants at 32-34 weeks gestation compared to $4539 for the moderate preterm group in this study (20). A local study by Cheah et al reported total cost per infant ranging from $26 to $3818 for babies between 1000-1500kg (1999 data and cost values) (11). In this study for the very preterm group, the total cost per infant ranged from MYR 3648 – MYR 48,943 ($994-$13,336) and cost per day ranged from MYR 376– MYR 782 ($102-$213). Differences in cost observed above may be due to factors such as time, geography, study design and sample, inflation rates, health financing system and hospital charge policies. In this study mean total admission cost per infant for extreme preterm intensive care was five times more than late preterm intensive care and fifteen times more than preterm minimal care. Geitona et al found a more conservative one and a half time increase in cost from moderate and late preterm to extreme preterm (15). Other studies found exponential rise in cost with reducing gestation and birth weight. Russell et al found the cost to be four times more for extreme preterm infants compared to the average preterm (8). Narang et al found the total admission cost for preterm infants less than 1000g was 4 times more than those weighing 1250g-1500g (17). The cost of neonatal care for infants below 1000 g was found to be 75% higher compared to those between 1000g to 1499g and more than four times higher than those weighing 1500g and more (3, 21). These findings relate to this study where the total cost of intensive care per infant for extreme preterm (mean birth weight 0.92kg) was more than one and a half times higher than very preterm group (mean birth weight 1.26kg), two and a half times higher than moderate preterm (mean birth weight 1.69kg) and five times higher than late preterm (mean birth weight 2.08kg). However it has been observed that more than two thirds of total preterm hospitalization cost belonged to infants who were not extremely preterm (22).

Gender, birth weight and length of stay as cost predictors in this study are supported by previous evidences. A compilation of related studies have shown that male preterm infants face higher risk of mortality and morbidity which contribute to higher cost (23). Morbidities include blindness, deafness and neurological disorders such as learning problems and cerebral palsy. Higher risks for complications of respiratory distress syndrome and intra-ventricular haemorrhage occur for those born at 23-32 weeks of gestation and is more pronounced in the extreme preterm group (24, 25). In the acute stage of illness more preterm males need mechanical ventilation, inotropic support and require more surfactant (26). During convalescence more preterm males develop broncho-pulmonary dysplasia and require supplementary oxygen upon discharge (27). Better outcome in female preterm infants may be explained by faster maturation during gestation leading to more developed lungs and other organs that avoids complications (23, 25). Furthermore in the event of hypoxic stress during labour preterm females have significantly higher catecholamine levels than males as a protective response. A comparison between male and female infants who received intensive care in this study is shown in Table 5. Despite similarities in mean gestational age and birth weight male infants generally required more resources and cost, reflecting findings in the aforementioned studies.

**Table 5:** Gender comparison (intensive care)

|  | Mean (range) |  |
| --- | --- | --- |
|  | Male (n=43) | Female (n=50) |
| Gestation (weeks) | 31<br>(26-36) | 31<br>(25-36) |
| Birth weight (kg) | 1.55<br>(0.61-2.73) | 1.52<br>(0.79-2.69) |
| Resource utilization |  |  |
| Length of stay (days) | 43<br>(2-167) | 41<br>(6-111) |
| Ventilation (days) | 17<br>(0-104) | 12<br>(0-56) |
| Parenteral nutrition (units) | 17<br>(0-105) | 10<br>(0-38) |
| Mobile imaging (units) | 6<br>(1-23) | 5<br>(0-18) |
| Transfusion (units) | 3<br>(0-20) | 1<br>(0-10) |
| Cost |  |  |
| Consumables (MYR) | 5009<br>(104-31,961) | 3140<br>(12-12,629) |
| Admission (MYR) | 19303<br>(534-98,649) | 15959<br>(374-49,356) |

Birth weight as a predictive factor for admission cost is consistent with findings by Akman et al which reported birth weight as among powerful predictive factors for hospital costs (19). Cost analysis studies too reflected changes of cost in (inverse) relation to birth weight and this includes a local study by Cheah et al. which reported that the total cost per infant ranged from $26 to $3818 for babies between 1000-1500kg (1999 data and cost values) (11). Narang et al found the total admission cost for preterm infants less than 1000g was 4 times more than those weighing 1250g-1500g (17). Cost of neonatal care for infants below 1000 g was found to be 75% higher compared to those between 1000g to 1499g and more than four times higher than those weighing 1500g and more (3, 21). In summary increasing prematurity either by gestational age or birth weight is associated with exponential increase in hospital cost (9). Length of stay as one of the predictive factors of total admission cost is consistent with findings in a study by Moran et al. which looked at total cost for survivors and non-survivors in adult intensive care (28). Richardson et al advocated reducing length of stay to reduce NICU costs due to high cost per diem (14). However it was emphasized that different clinical management aims between term and preterm infants required separate strategies between the two groups to reduce length of stay. Term infants require admission only in the event of an illness and length of stay would depend on its severity and the effectiveness of treatment. In contrast preterm infants require admission (often prolonged) due to inherent complications and to achieve maturity and ideal weight before discharge. Nevertheless, utilizing length of stay as a sole cost-reducing measure has its pitfall. Evans et al. and Kerlin & Cooke stressed that reducing length of stay will not have significant effect on reducing hospital costs (29, 30). This was based on the argument that variable costs comprised only about 20% of an intensive care unit (ICU) admission cost and attempts to shorten the duration of stay may not have any appreciative impact. In contrast ICU policy changes which affect fixed costs will have larger bearing upon total expenditure. Furthermore ICU cost is highest during the initial period of admission which corresponds with the intensity of care and denies reducing length of stay as a cost-saving strategy. DeRienzo et al. used an NICU simulation model and found that reducing length of stay did not uniformly reduce hospital resource utilization or hospital cost (31). It demonstrated that longer length of stay literally improved clinical outcomes and reduced cost thus suggesting that emphasis on clinical outcomes must accompany efforts to reduce length of stay.

The analysis by degree of prematurity in this study provided in-depth understanding of differences in resource consumption and ultimately cost of admission. Among the important findings were mean admission cost per infant increased with level of care and prematurity, more than two thirds of total admission cost per preterm was attributable to NICU specific overhead (fixed) costs and personnel salary and laboratory investigations were the cost drivers for overhead and consumables (variable) costs respectively. Meanwhile gender, birth weight and length of stay were the cost predictors whereby total admission cost was higher for male infants, lower birth weight and longer length of stay. The fact that preterm infants account for the bulk of NICU admissions make these findings and economic evaluation altogether relevant to guide policy and decision making for neonatal care in Malaysia. An example would be policies and measures to optimize quantity and quality of human resource as personnel salary was found to contribute three quarters of NICU specific overhead cost in this study. Thus economic evaluation should be widely applied as it is integral to any plans for establishment, running or expansion of services especially with the advent of new and costly treatment modalities. Prior to this study there had been only one economic evaluation performed which looked at cost effectiveness of NICU care in Malaysia for infants between 1000 and 1500 g birth weight (11). To the best of our knowledge this is the first study in the region to detail the impact of increasing prematurity on provider cost and also develop a cost prediction equation. Utilizing results from this study can facilitate cost-effectiveness evaluation of current and new management strategies in all preterm categories.

There were limitations in this study. Preterm deaths were excluded thus costs involved in management of this group of patients were not accounted for. Complications that occurred in each preterm infant recruited were not taken into account thus missing the diagnosis-specific cost analysis. Nevertheless, the cost of management for the complications would had been included as the length of stay would have been prolonged. Several key strengths too can be highlighted. Findings from provider cost analysis are generalizable for preterm care in Malaysia as hospitals under the Ministry of Health have similar administrative, financial and organizational structures and NICU clinical management follow common practice guidelines. This study had employed a mixed method of top down and bottom up costing for care provider cost. By employing bottom up micro costing for the variable (consumables) component a more refined cost estimate was obtained. This method provided a balance between less accurate but easier to perform gross costing and the more accurate but resource consuming micro costing. Costs were analysed by degree of prematurity making it the first local study to provide a comprehensive data on the impact of increasing prematurity on provider cost. This study produced the first cost prediction of preterm NICU admission for Malaysia.

## Acknowledgements

The authors would like to acknowledge consultant pediatricians Dr. Thiyagar Nadarajaw, Dr. Tan Ying Beih and Dr. Choo Chong Ming for their willingness to participate in this study. Special mention to NICU doctors and nurses of both study centers who diligently assisted in data collection and analysis. We would also like to express our gratitude to the Director General of Health, Malaysia for permission to conduct and publish this study.

## Author contributions

Conceptualization - Hadzri Zainal, Maznah Dahlui, Tin Tin Su
Data Curation – Hadzri Zainal
Formal Analysis - Hadzri Zainal, Maznah Dahlui, Tin Tin Su, Shahrul Aiman Soelar
Funding Acquisition - Hadzri Zainal, Maznah Dahlui, Tin Tin Su
Investigation - Hadzri Zainal
Methodology - Hadzri Zainal, Maznah Dahlui, Tin Tin Su, Shahrul Aiman Soelar
Project Administration - Hadzri Zainal
Resources - Hadzri Zainal
Software - Hadzri Zainal, Shahrul Aiman Soelar
Supervision - Maznah Dahlui, Tin Tin Su
Validation - Hadzri Zainal, Maznah Dahlui, Tin Tin Su
Visualization - Hadzri Zainal
Writing – Original Draft: Hadzri Zainal
Writing – Review & Editing: Hadzri Zainal, Maznah Dahlui, Tin Tin Su, Shahrul Aiman Soelar

